# Structural insights on TRPV5 gating by endogenous modulators

**DOI:** 10.1101/338798

**Authors:** Taylor E.T. Hughes, Ruth A. Pumroy, Aysenur Torun Yazici, Marina A. Kasimova, Edwin C. Fluck, Kevin W. Huynh, Amrita Samanta, Sudheer Mulugu, Z. Hong Zhou, Vincenzo Carnevale, Tibor Rohacs, Vera Y. Moiseenkova-Bell

## Abstract

TRPV5 is a transient receptor potential channel involved in calcium reabsorption. Here we investigate the interaction of two endogenous modulators with TRPV5. Both phosphatidylinositol 4,5-bisphosphate (PI(4,5)P2) and calmodulin (CaM) have been shown to directly bind to TRPV5 and activate or inactivate the channel, respectively. Using cryo-electron microscopy (cryo-EM), we determined TRPV5 structures in the presence of dioctanoyl PI(4,5)P2 and CaM. The PI(4,5)P2 structure revealed a novel binding site between the N-linker, S4-S5 linker and S6 helix of TRPV5. These interactions with PI(4,5)P2 induce conformational rearrangements in the lower gate, opening the channel. The CaM structure revealed two TRPV5 C-terminal peptides anchoring a single CaM molecule and that calcium inhibition is mediated through a cation-π interaction between Lys116 on the C-lobe of calcium-activated CaM and Trp583 at the intracellular gate of TRPV5. Overall, this investigation provides insight into the endogenous modulation of TRPV5, which has the potential to guide drug discovery.

## INTRODUCTION

The calcium ion is vital for an array of cellular functions, and in the human body the kidneys regulate calcium homeostasis by filtration and reabsorption^1^. Around 99% of calcium is reabsorbed by the kidney tubules^1^, with ~15% of that reabsorption occurring via transient receptor potential vanilloid 5 (TRPV5) channels in the distal convoluted tubule and collecting tubule. TRPV5 is specialized for this task and is only expressed in the apical membrane of kidney epithelial cells in the distal convoluted tubule and collecting tubule ^2,3^ When open, TRPV5 allows calcium in the urine to flow along a concentration gradient through the channel pore into the cell. This gradient is maintained by the calcium sequestering protein calbindin, which delivers calcium to active transport proteins at the basolateral membrane of the epithelium, which then export the ions into the blood stream^2,3^. Though this mechanism of calcium reabsorption is responsible for only ~15% of total calcium reabsorbed in the kidney, TRPV5 activity is critical for the homeostatic balance of calcium^2,3^. This is exemplified by TRPV5 knock out mice, which have been reported to have systemic calcium imbalance in the form of hypercalciuria and bone mineral loss^2,3^. In humans, single nucleotide polymorphisms in TRPV5 in African-American populations result in a significant increase in calcium reabsorption that is correlated with a lower risk of nephrolithiasis^2,3^. Together, these observations suggest that TRPV5 could be a potential drug target for human disorders involving altered calcium homeostasis.

TRPV5 is a calcium selective channel that displays constitutive activity in the presence of basal levels of the membrane phospholipid phosphatidylinositol 4,5-bisphosphate (PI(4,5)P_2_)^2^. The TRPV5 channel is tetrameric and consists of classic TRPV family features, including 6 transmembrane helices (S1-S6), N-terminal ankyrin repeats (ARD), and the TRP domain^4^ (Fig. 1A). TRPV6 is a closely related epithelial Ca^2+^ channel that shows a high level of sequence homology with TRPV5^3^; these two channels have a much lower level of sequence homology with other members of the TRPV subfamily. The regulation and biophysical properties of TRPV5 and TRPV6 are similar, and they are functionally quite different from the rest of the TRPV subfamily^3^. PI(4,5)P_2_ is found in the inner leaflet of the plasma membrane and it has been shown to be essential for the activity of both TRPV5^5,6^ and TRPV6^7^. For TRPV6, the activating effect of PI(4,5)P_2_ was demonstrated in planar lipid bilayers, providing definitive evidence for direct effect on the channel^8^. However, the molecular details of the PI(4,5)P_2_ interaction with TRPV5 and TRPV6 are essentially unknown.

**Figure 1.**
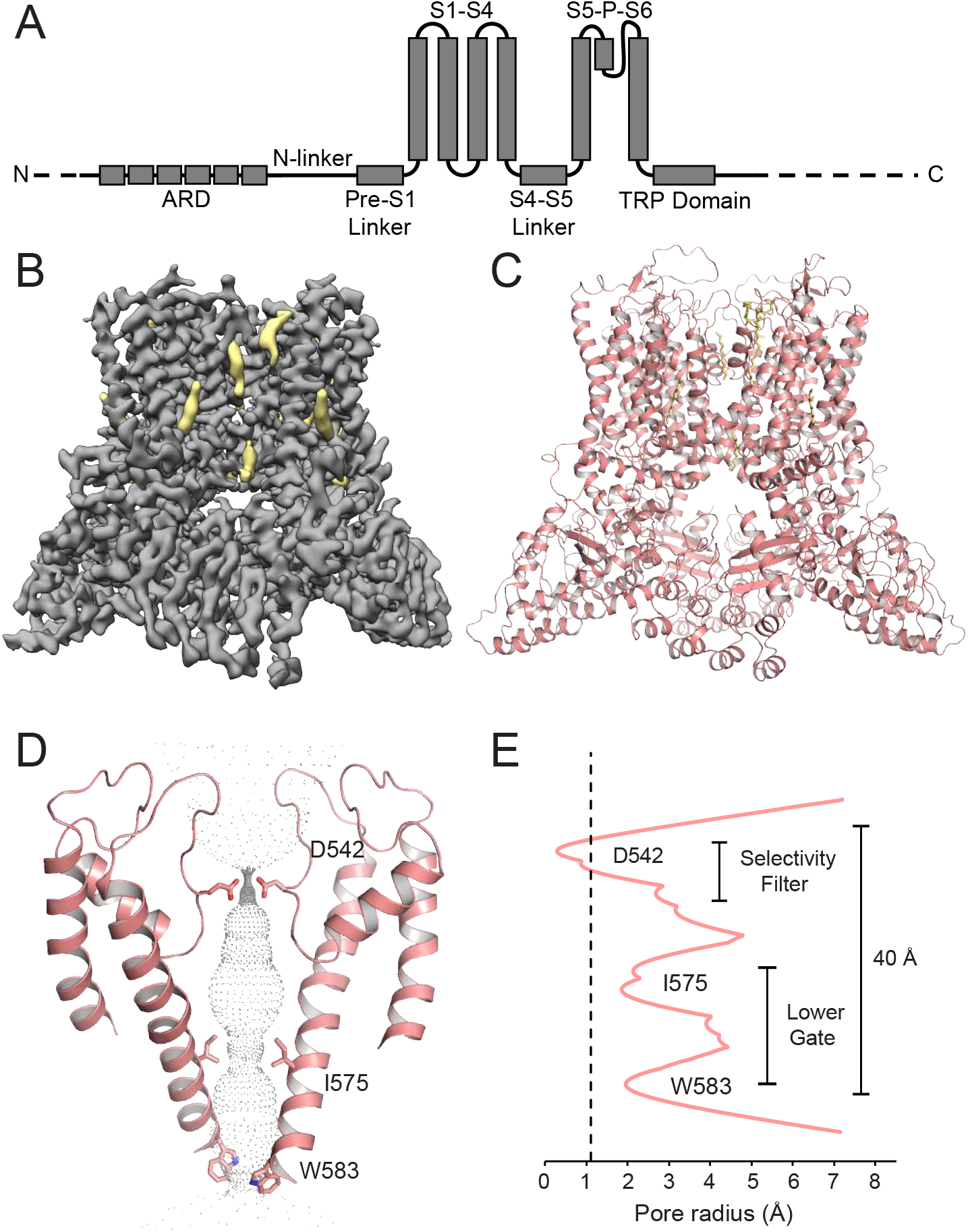
Lipid-bound TRPV5 structure in detergent. (A) A schematic representation of the TRPV5 domains per channel monomer. Dashed lines indicate regions for which a model could not be built. (B) Density map of lipid-bound TRPV5 at 3.9Å resolution. Density for TRPV5 is shown in grey and the densities attributed to annular lipids are shown in khaki. (C) Cartoon representation of the lipid-bound TRPV5 model. The TRPV5 tetramer is depicted as pink cartoons and annular lipids as khaki sticks. (D) Cartoon representation of the lipid-bound TRPV5 pore highlighting the three constriction points in the selectivity filter and lower gate. (E) Plot of pore radii of lipid-bound TRPV5 as a function of distance through the pore. The dotted line indicates the radius of a dehydrated calcium ion.

To prevent excess flow of calcium ions into the cell, both TRPV5 and TRPV6 have been shown to rapidly inactivate through a calcium-dependent mechanism^9^. Calmodulin (CaM), a calcium sensing protein, has been shown to be involved in this inactivation by directly interacting with the last thirty amino acids of the TRPV5 C-terminus^10^. While the role of the distal C-terminal binding site is well established in CaM-mediated inactivation of TRPV5^10^ and the closely related TRPV6^11–13^, the binding stoichiometry and the conformational changes that take place as a consequence of the binding of CaM to the channel have been unclear^3^. The absence of this information prevents us from further understanding TRPV5 function and regulation in the kidney.

In this study, we used cryo-electron microscopy (cryo-EM) to uncover the molecular mechanisms of TRPV5 gating via endogenous modulators. Investigating TRPV5 modulation with cryo-EM permitted us to gain insight regarding TRPV5 gating and potentially form the basis for rational drug design for the treatment and prevention of hypercalciuria and nephrolithiasis in future studies.

## RESULTS

### Structures of TRPV5 in the lipid-bound and in the PI(4,5)P_2_-bound states

TRPV5 is part of the tightly regulated system of calcium homeostasis in the human body^2^. Two endogenous mechanisms of regulation, binding of PI(4,5)P_2_ or CaM to TRPV5, result in activation or inactivation of the channel^13^, respectively. To understand these mechanisms of modulation, first we used cryo-EM to determine the structure of detergent solubilized full-length rabbit TRPV5 in the presence of the 200 μM dioctanoyl (diC_8_) PI(4,5)P_2_, a soluble form of PI(4,5)P_2_. This concentration is ~3 times the EC50 of diC_8_ PI(4,5)P_2_ for TRPV6, and TRPV5 has a slightly higher apparent affinity for diC_8_ PI(4,5)P_2_^14^, thus this concentration was expected to result in near saturation of TRPV5 with diC_8_ PI(4,5)P_2_. During the cryo-EM structure determination process, one stable class emerged and was refined to an overall resolution of 3.9 Å (Fig. 1B, Sup. Fig. 1, Sup. Fig. 2). The vast majority of this TRPV5 cryo-EM map is at high resolution (3-3.5 Å) as side chains are clearly visible in the transmembrane region of the channel (Sup. Fig. 1, Sup. Fig. 3). This allowed for accurate model building in both the transmembrane domain (TMD) and the ankyrin repeat domain (ARD) (Fig. 1C, Sup. Fig. 3). Highly flexible areas such as the very distal C- and N-termini were unable to be resolved. Interestingly, five non-protein densities per monomer were identified in this cryo-EM map (Fig. 1B). None of these could be identified as diC_8_ PI(4,5)P_2_, and were attributed to annular lipids that had high enough affinity for TRPV5 to be co-purified with the protein. Similarly shaped non-protein densities were also assigned as lipids in a recently reported human TRPV6 cryo-EM structure^15^. Therefore, we will refer to this channel reconstruction as lipid-bound TRPV5.

The lipid-bound TRPV5 pore contains three residues that are involved in pore constriction: Asp542, Ile575 and Trp583 (Fig. 1D). Unlike the previously published structure of TRPV5^4^, there is clear density in the lipid-bound TRPV5 map indicating that the four Asp542 that constitute the highly specific Ca^2+^ selectivity filter are pointing directly into the pore in a similar fashion to the rat TRPV6 crystal structures^16,17^. The lipid-bound TRPV5 lower gate appears to consist of Ile575 and Trp583 (Fig. 1D-E). These residues do not constrict the pore to the point of blocking ion translocation, indicating that the lower gate is open (Fig. 1E), as seen in the human TRPV6 cryo-EM structure^15^. This suggests that the pore of the lipid-bound TRPV5 structure may be in a closed pre-open conformation.

Based on these findings, we decided to increase diC_8_ PI(4,5)P_2_ concentration to 400 μM and use nanodiscs to capture TRPV5 in the PI(4,5)P_2_-bound state. During the cryo-EM structure determination process we noted that even before masking and 3D classification in RELION^18,19^ the initial 3D reconstruction contained a nonprotein density that resembled diC_8_ PI(4,5)P_2_ in both size and shape (Fig. 2A, Sup. Fig. 4). Next, we used particle subtraction and 3D classification that was focused on the diC_8_ PI(4,5)P_2_ binding site using a mask which included the entire TRP domain helix, the base of the S6 helix, and the head group of the diC_8_ PI(4,5)P_2_ (Sup. Fig. 4). This classification yielded three classes with good features: one without diC_8_ PI(4,5)P_2_ and two with diC_8_ PI(4,5)P_2_ (Sup. Fig. 4). The class without diC_8_ PI(4,5)P_2_ yielded a TRPV5 structure at 4.4Å, which clearly resembled the lipid-bound TRPV5 in detergent (Sup. Fig. 5). Due to high similarity between the detergent and nanodisc lipid-bound TRPV5 structures (Sup. Fig. 5), we will only describe the lipid-bound TRPV5 in detergent as it was determined at a higher resolution (Sup. Fig. 1, Sup. Fig. 3, Sup. Fig. 5). The two classes with diC_8_ PI(4,5)P_2_ density, after additional classification and refinement, produced a single TRPV5 structure in a diC_8_ PI(4,5)P_2_-bound state at 4.0 Å (Fig. 2B, Sup. Fig. 6, Sup. Fig. 7). All structures identified in nanodiscs revealed non-protein densities in similar positions to those attributed to annular lipids in the lipid-bound TRPV5 structure in detergent (Fig. 1B, Fig. 2A-B, Sup. Fig. 5).

**Figure 2.**
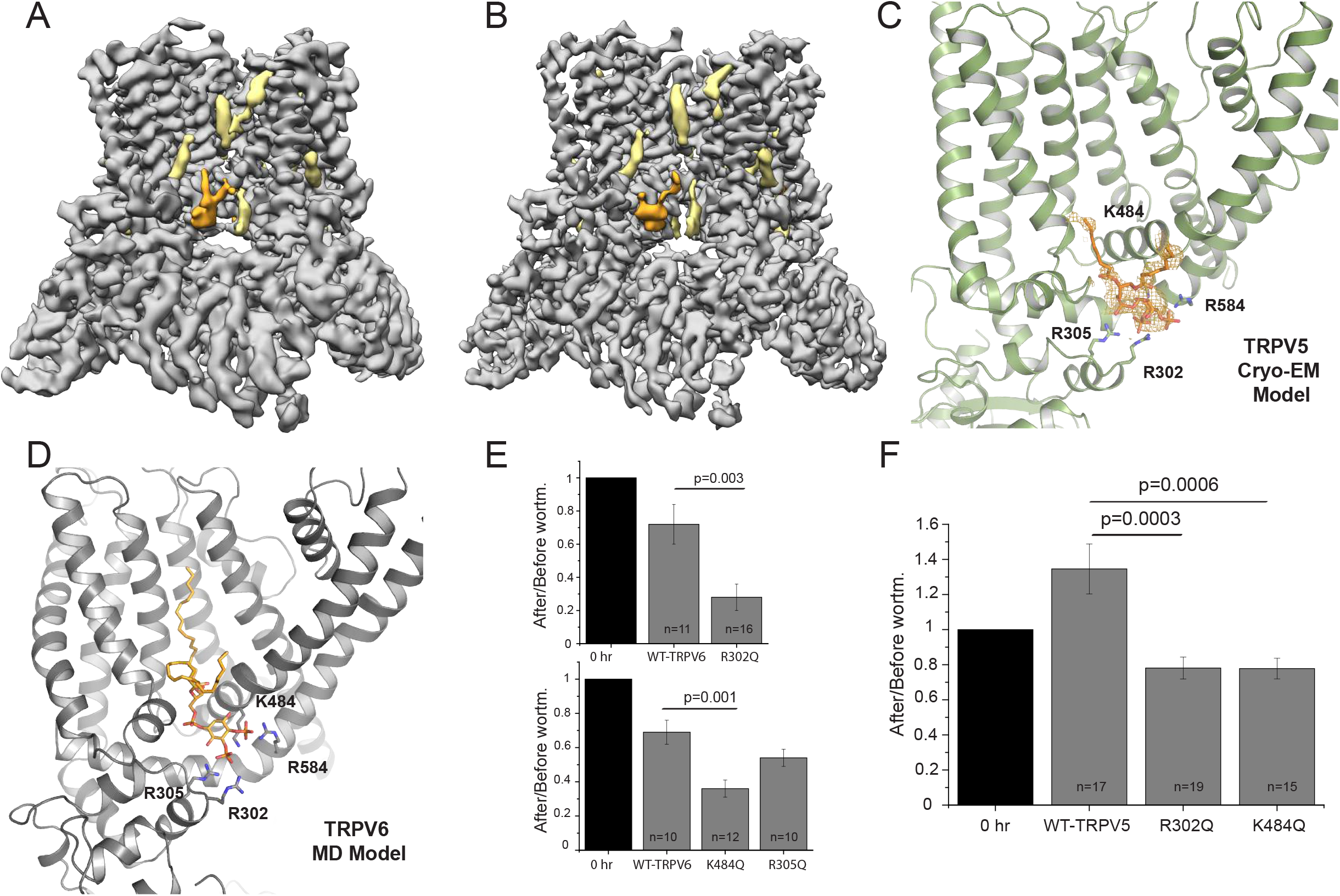
Phosphatidylinositol 4,5-bisphosphate (PI(4,5)P2)-bound TRPV5 structure and PI(4,5)P2-bound TRPV6 modelling. (A) The initial reconstruction of PI(4,5)P2-bound TRPV5 in nanodiscs before masking and 3D classification. TRPV5 density is shown in grey, annular lipids are shown in khaki, and PI(4,5)P2 is shown in orange. (B) PI(4,5)P2-bound TRPV5 cryo-EM density in nanodiscs after focused 3D classification. TRPV5 density is shown in grey, annular lipids are shown in khaki, and PI(4,5)P2 is shown in orange. (C) Zoomed-in view of the TRPV5 PI(4,5)P2 -binding pocket. The PI(4,5)P2 binding site in the TRPV5 channel is located between the N-linker (R302, R305), S4-S5 linker (K484) and the S6 helix (R584) of the channel. (D) A model produced by molecular dynamics (MD) simulations of the predicted interaction between the homologous TRPV6 channel and PI(4,5)P2. (E) Neutralization of key PI(4,5)P2 interacting residues increases sensitivity to depletion of PI(4,5)P2 in TRPV6. The TRPV6 currents in oocytes measured before and after incubation with 35 μM wortmannin for 1 hr. Current values after wortmannin treatment are normalized to current values before treatment. P-values for significance values are shown after rounding to the first non-zero digit (analysis of variance). (F) Neutralization of key PI(4,5)P2 interacting residues increases sensitivity to depletion of PI(4,5)P2 in TRPV5. The TRPV5 currents in oocytes measured before and after incubation with 35 μM wortmannin for 1 hr. Current values after wortmannin treatment are normalized to current values before treatment. P-values for significance values are shown after rounding to the first non-zero digit (analysis of variance).

The PI(4,5)P_2_ binding site in the TRPV5 channel is located between the N-linker (Arg302, Arg305), S4-S5 linker (Lys484) and the S6 helix (Arg584) of the channel (Fig. 2C). These results show an excellent correlation with predictions by earlier molecular dynamics (MD) simulations performed on the homologous TRPV6 channel (Fig. 2D, Sup. Fig. 8). These simulations predicted that residues Arg302, Arg305, and Lys484 are involved in the TRPV6 interaction with PI(4,5)P_2_ (Fig. 2D, Sup. Fig. 8). Moreover, when these predicted PI(4,5)P_2_ interacting TRPV6 residues (Arg302, Arg305, Lys484) were mutated to glutamine, the channel had increased sensitivity to depletion of PI(4,5)P_2_ with high concentrations of wortmannin, which inhibits PI4-kinases (Fig. 2E). This result indicates reduced apparent affinity for PI(4,5)P_2_, which is expected if a PI(4,5)P_2_ interacting residue is mutated^20^. The current amplitudes of the Arg302Gln and Lys484Gln mutants were also significantly lower than wild type TRPV6 (Sup. Fig. 9), which is also consistent with decreased apparent affinity of the channel for PI(4,5)P_2_. The Arg305 residue showed a less clear contact with PI(4,5)P_2_ compared to Arg302 and Lys484, interacting with only 3 out of 4 subunits in MD simulations (Sup. Fig. 8). Accordingly, this mutant showed a less pronounced trend to be inhibited more by wortmannin and had a smaller amplitude, but these effects did not reach statistical significance from wild type (Fig. 2E, Sup. Fig. 9). When we mutated residues Arg302 and Lys484 to glutamine in TRPV5 and performed analogous experiments, we observed that the TRPV5 channel also had significantly increased sensitivity to the depletion of PI(4,5)P_2_ with high concentrations of wortmannin, suggesting that both TRPV5 and TRPV6 share a similar PI(4,5)P_2_ binding site (Fig. 2F). Unlike for TRPV6, current amplitudes for these mutants were not statistically significantly different from wild type TRPV5 (Sup. Fig. 9).

Based on the structure of PI(4,5)P_2_ -bound TRPV5 compared to the lipid-bound TRPV5, the binding of diC_8_ PI(4,5)P_2_ to TRPV5 appears to induce conformational changes related to channel activity. The pore of the channel widened, which could allow for the flow of hydrated Ca^2+^ ions (Fig.3A-D). Specifically, the Asp542 residues, which line the selectivity filter of the channel and coordinate Ca^2+^ ions, point upwards in the extracellular vestibule of the channel instead of towards the ion conduction pathway as seen in the lipid-bound structure (Fig. 3D). The distance between the center of the carboxylate oxygen atoms of the Asp542 residues in this structure is ~3 Å, which would allow for partially hydrated Ca^2+^ ion to flow though the pore of the channel (Fig. 3C-D). The lower gate of the channel is also wide open (~12 Å compared to ~8 Å in the lipid-bound state) due to the movement of the S6 helix and Trp583 upon binding of diC_8_ PI(4,5)P_2_ (Fig. 3E), which would facilitate the movement of partially hydrated Ca^2+^ ions through the pore (Fig. 3C). Additionally, diC_8_ PI(4,5)P_2_ binding caused global conformational changes in the S6 and TRP helices as well as the S4-S5 linker when compared to the lipid-bound structure (Fig. 3F).

**Figure 3.**
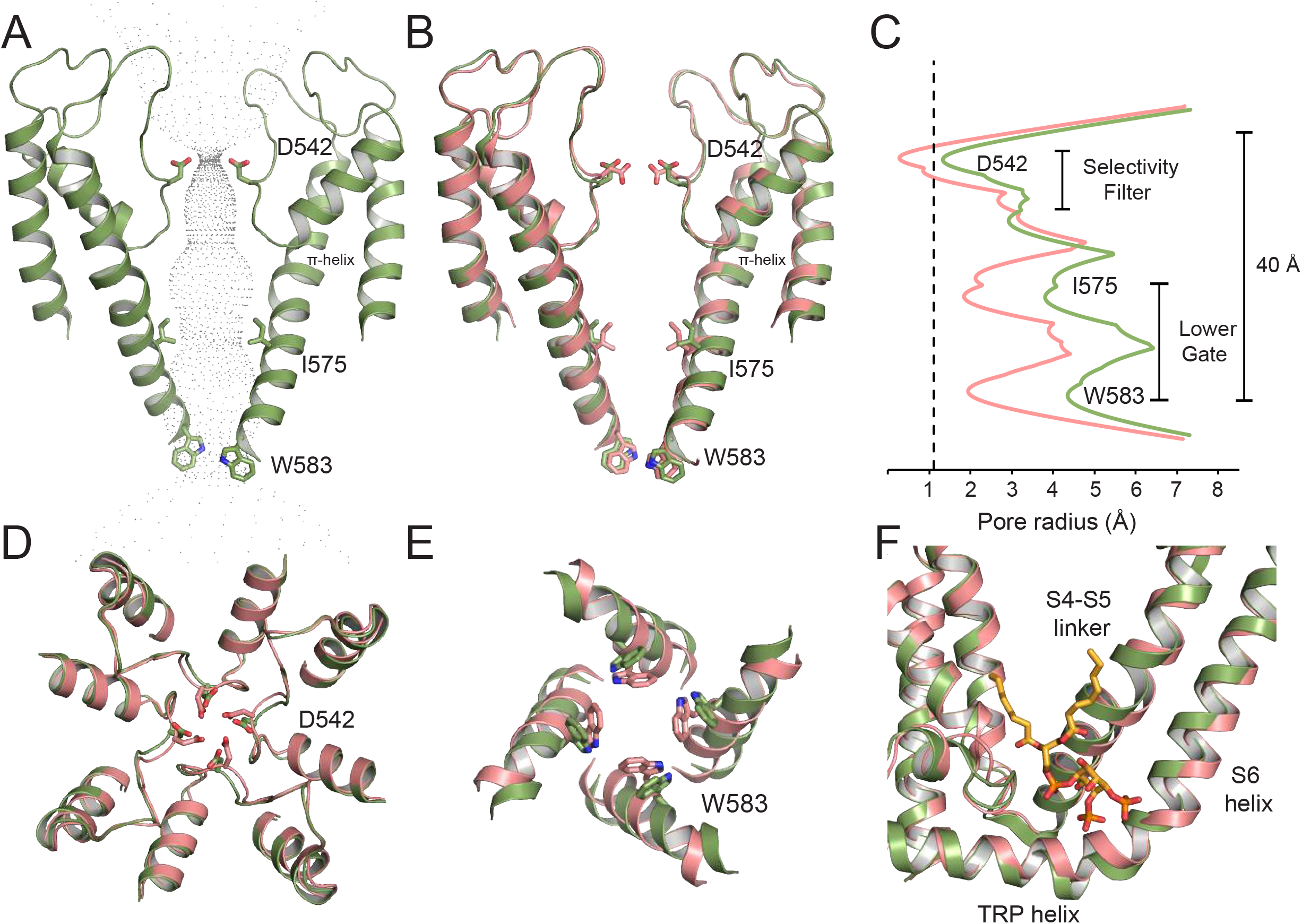
Comparison between lipid-bound and PI(4,5)P2-bound TRPV5. (A) The pore diagram of PI(4,5)P2-bound TRPV5 is shown in green. (B) Lipid-bound TRPV5 (pink) superimposed onto the PI(4,5)P2-bound (green) pore diagram shows a slight shift in the S6 helices and important gating residues originating at the π-helix. (C) Plot of pore radii of lipid-bound TRPV5 (pink) and PI(4,5)P2-bound TRPV5 (green) as a function of distance through the pore. The dotted line indicates the radius of a calcium ion. (D) Extracellular view of the tetrameric selectivity filter of lipid-bound (pink) and PI(4,5)P2-bound TRPV5 (green). (E) Intracellular view of the tetrameric lower gate of lipid-bound (pink) and PI(4,5)P2-bound TRPV5 (green). (F) Zoomed in view of the overlayed TMDs of lipid-bound (pink) and PI(4,5)P2-bound TRPV5 (green). DiC8 PI(4,5)P2 is shown in orange sticks.

Comparison between the diC_8_ PI(4,5)P_2_-bound and lipid-bound TRPV5 structures revealed details of the conformational changes induced by PI(4,5)P_2_ binding (Fig. 4). The inositol ring of the diC_8_ PI(4,5)P_2_ head group is positioned close to the lower gate of the channel, where phosphate groups at position four and five can interact directly with Arg584 and Arg302, respectively (Fig. 4A and 4C). Specifically, the phosphate group at the position four forms a salt bridge with the Arg584 on the S6 helix. This interaction is facilitated by the rotation of the Arg584 towards the PI(4,5)P_2_, which pulls the S6 helix away from the center of the pore, starting at the π-helix (Fig. 3B), and causes an outward shift of the S4-S5 linker (Fig. 3F). The rotation of Arg584 also induces the lengthening of the S6 helix and shortens the TRP helix (Fig. 3F, Fig. 4A-B): in the lipid-bound TRPV5 structure, the transition between the S6 helix and the TRP helix occurs between Arg584 and Val585, while in the diC_8_ PI(4,5)P_2_-bound TRPV5 structure this transition occurs between residues Gln587 and Glu588. The extension and rotation of the S6 helix pulls Trp583 out of the pore and stabilizes its position through an interaction with Gln587, thus opening the lower gate. The shortening of the TRP helix causes Glu588 to pull away from Arg302, allowing Arg302 to interact with the phosphate group at position five of the diC_8_ PI(4,5)P_2_ head group (Fig. 4C-D). Additionally, Lys484 interacts with the diC_8_ PI(4,5)P_2_ inositol ring, which further electrostatically stabilizes the position of diC_8_ PI(4,5)P_2_ in its binding pocket (Fig. 4C-D, Sup. Movie 1).

**Figure 4.**
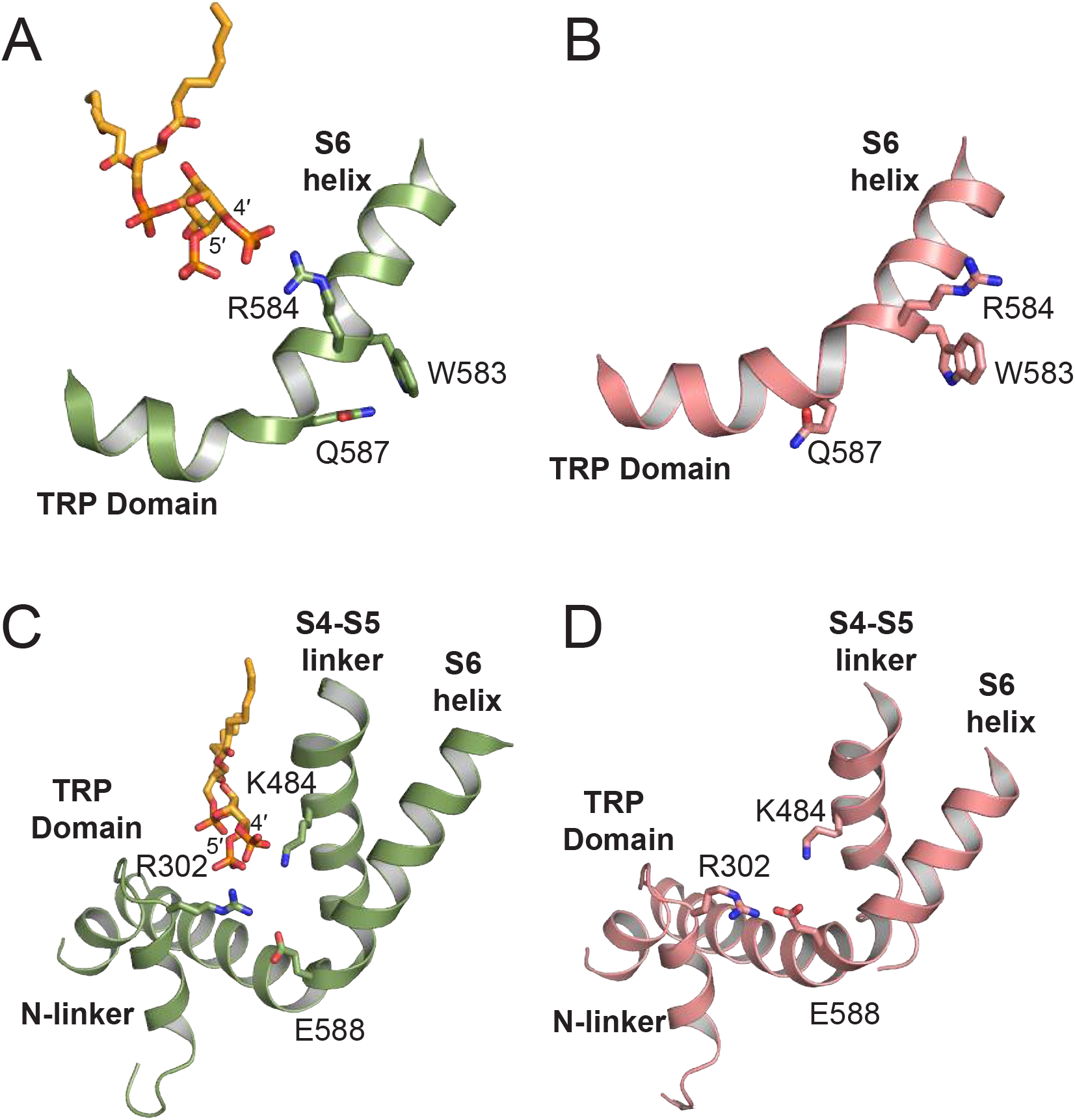
Interactions of TRPV5 with PI(4,5)P2. (A) View of the lower S6 helix and TRP domain of PI(4,5)P2-bound TRPV5. The PI(4,5)P2-bound TRPV5 is shown in green and the PI(4,5)P2 in orange. (B) View of the lower S6 helix and TRP domain of the lipid-bound TRPV5 structure (pink). (C) The 5’ phosphate of PI(4,5)P2 (orange) interacts with R302 of the N-linker and K484 of the S4-S5 linker in the PI(4,5)P2-bound TRPV5 structure. (D) The lipid-bound structure showing the N-linker, S4-S5 linker, S6 helix and TRP domain in pink.

### TRPV5 inhibition by calmodulin

In order to investigate the mechanism of TRPV5 inactivation by CaM, we incubated detergent solubilized full-length rabbit TRPV5 with calcium-activated rat CaM in a 1:20 molar ratio in the presence of 10 mM CaCl_2_. The stoichiometry of TRPV5 to CaM has been speculated to be between one and four molecules of CaM per tetramer^3,6,21–23^ and the affinity between CaM and the TRPV5 C-terminus estimated at ~0.3 μM^24^, therefore the high molar ratio was used to ensure TRPV5 saturation with CaM. This sample yielded a cryo-EM map at 4.4 Å without applied symmetry (Fig. 5A, Sup. Fig. 10, Sup. Fig. 11, Sup. Fig. 12). In this structure, a single CaM molecule is bound to the intracellular section of TRPV5 at the base of the pore (Fig. 5A). Both lobes of CaM are resolved and each bound to different sections of the TRPV5 C-terminus (Fig. 5A-B, Sup. Fig. 12). Despite the interaction with CaM, the CaM-bound conformation of TRPV5 is almost identical to the lipid-bound conformation of TRPV5, with an RMSD of ~0.6A overall and a very similar pore profile (Fig. 5C-D). Additionally, annular lipids were also identified in the CaM-bound TRPV5 structure in similar positions to those in the lipid-bound TRPV5 structure in detergent (Fig. 1B, Fig. 5A).

**Figure 5.**
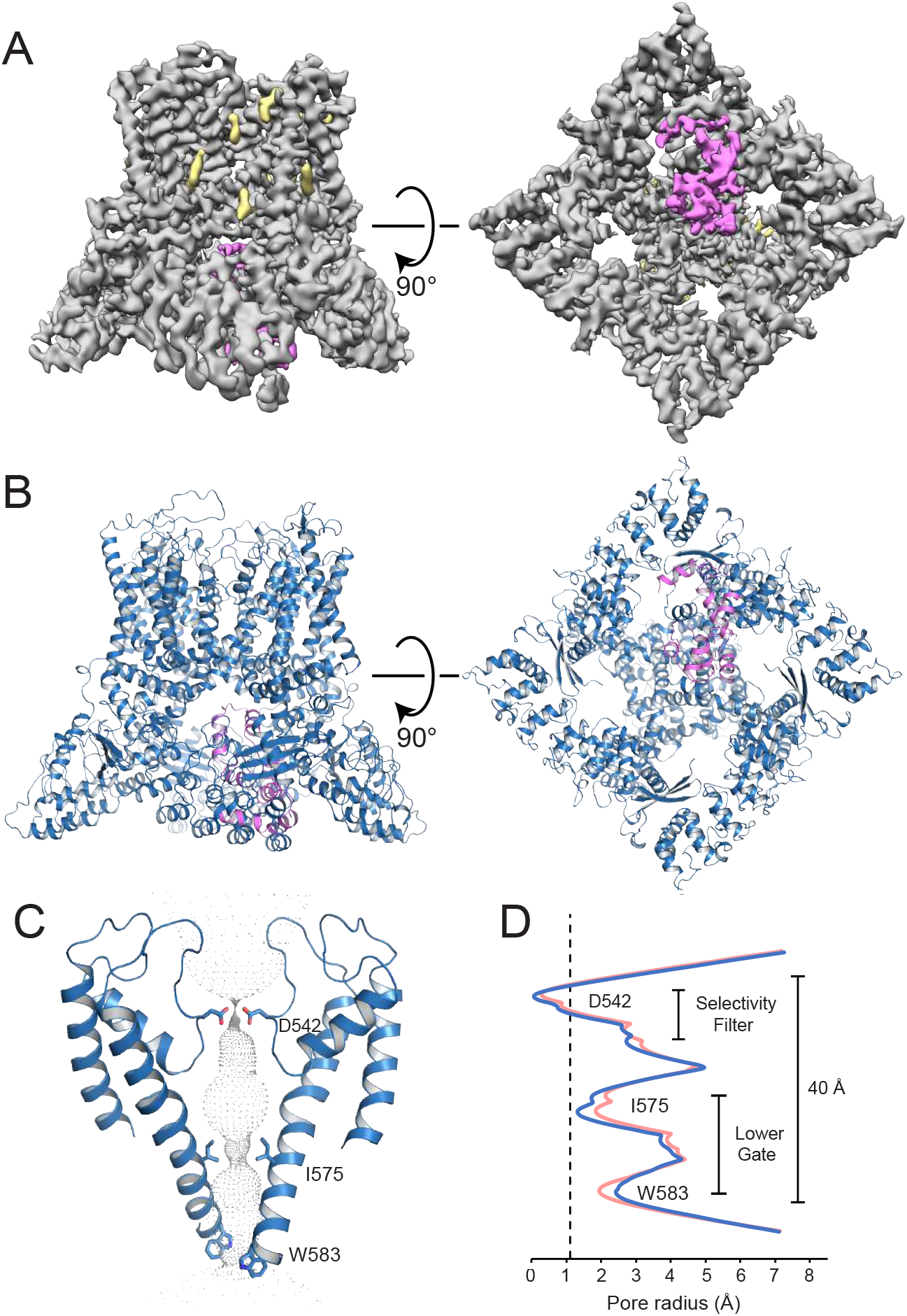
CaM-bound structure of TRPV5. (A) Side and bottom view of the CaM-bound TRPV5 density map at 4.4Å resolution. Density for TRPV5 is shown in grey, annular lipids shown in khaki, and CaM shown in hot pink. (B) Cartoon representation of the CaM-bound TRPV5 model in a side and bottom view. TRPV5 is shown in blue and CaM is shown in hot pink. (C) Pore diagram of CaM-bound TRPV5. Constriction residues are labeled and shown as sticks. (D) Plot of pore radii of CaM-bound TRPV5 as a function of distance through the pore is shown in blue. For reference, the lipid-bound TRPV5 pore graph is shown in pink. The dotted line indicates the radius of a calcium ion.

CaM is bound with the C-lobe at the base of the pore and the N-lobe resting against the ankyrin repeats of two adjacent TRPV5 monomers (Fig. 5A-B). A portion of the CaM N-lobe (Gly41 -Ile64) and the linker between the CaM two lobes (Lys78-Ser82) are visible only as broken density and were excluded from the model (Fig. 6A). A peptide from the C-terminus of TRPV5 (His699-Thr709) is bound to the CaM C-lobe, with TRPV5 Trp702 making critical contacts with the CaM binding pocket, as previously reported^23^ (Fig. 6A and 6C). This peptide was arbitrarily assigned to TRPV5 chain A in the model, but no connections could be seen between the density for this peptide and any TRPV5 monomer. An extension of the TRPV5 chain A C-terminus (Asn640-Lys652) forms a helix and is bound to the hydrophobic binding pocket of the CaM N-lobe (Fig. 6B). This interaction is mediated by numerous hydrophobic contacts: Val644, Leu645, Val648, and Phe651 on the TRPV5 side and Phe20, Leu33, Val36, and Leu64 on the CaM side (Fig. 6A and 6D). A loop of the CaM C-lobe (Gly114-Thr118) crosses over the bottom of the TRPV5 pore, placing Lys116 directly into the pore. The ε-amino group of Lys116 sits in between the four Trp583 residues of the lower gate, forming a cation-π interaction and thereby blocking the flow of ions through the pore (Fig. 6E). To test the functional importance of Trp583 in TRPV5 inhibition by CaM, we performed excised inside out patch clamp experiments. Wild type TRPV5 currents were robustly inhibited by Ca^2+^-CaM (0.2 μM CaM, 3 μM free Ca^2+^) applied to the inner leaflet of the patch membrane (Fig. 6F and 6H), as previous reported for TRPV6^13^. On the other hand, the mutation of Trp583 to Leu in TRPV5 was essentially not inhibited by Ca^2+^-CaM (Fig. 6G and 6H). Thus, this cation-π interaction appears to be critical for CaM-induced inhibition of TRPV5 (Fig. 6F-H).

**Figure 6.**
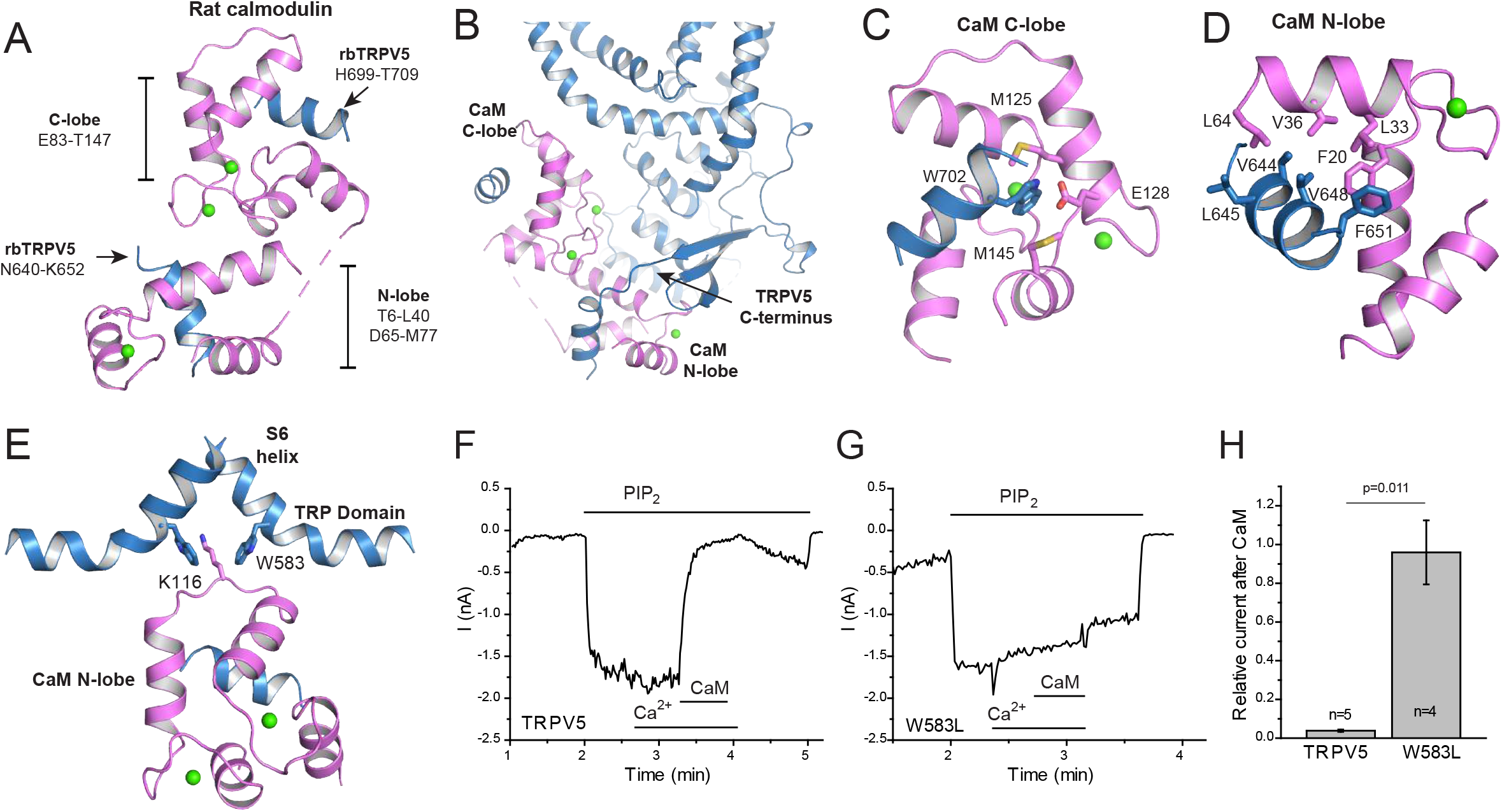
Mechanism of TRPV5 inactivation by CaM. (A) Structure of rat CaM bound to TRPV5 C-terminus fragments. Both the N-lobe and C-lobe were visualized along with N640-K652 and H699-T709 of TRPV5. CaM is shown in hot pink, and TRPV5 is shown in blue. Calcium ions bound to the N- and C-lobes of CaM are shown as green spheres. (B) CaM interaction with chain A of TRPV5. (C) CaM C-lobe interaction with W702 on the TRPV5 C-terminus. (D) CaM N-lobe makes multiple hydrophobic interactions along N640-K652. (E) The base of the TRPV5 pore is shown with K116 of CaM bound at W583, causing a steric blockage of calcium flow. (F) The excised inside out patch clamp experiments show that wild type TRPV5 is inactivated upon addition of Ca2^+^ activated CaM. Representative current trace is at -100 mV; the applications of 25 μM diC8 PI(4,5)P2, 0.2 μM CaM and 3 μM free Ca2^+^ are indicated by the horizontal lines. (G) The excised inside out patch clamp experiments show that Ca2^+^ activated CaM did not inactivate W583L TRPV5 mutant. Representative current trace is at -100 mV; the applications of 25 μM diC8 PI(4,5)P2, 0.2 μM CaM and 3 μM free Ca2^+^ are indicated by the horizontal lines. (H) Summary of the data, current amplitudes after CaM were divided by current values before CaM applications, p=0.011 (two-sample t-test).

## CONCLUSION

Here we have presented three structures of TRPV5, one in the presence of endogenous lipids that are likely to be involved in maintaining the structure of TRPV5, one bound to its endogenous activator, PI(4,5)P_2_ and one bound to its endogenous inhibitor, CaM. The high-resolution, lipid-bound TRPV5 has allowed for more accurate model building of TRPV5 than available previously^4^. This structure also identified the binding pockets for several high affinity lipids which were consistently seen in all TRPV5 structures presented here. These binding pockets have the potential to be druggable areas of the TRPV5 channel as they are both membrane and solvent accessible and are in contact with regions that have been reported to be able to transmit conformational rearrangements to the pore in other TRPV family channels^4,15,25^.

PI(4,5)P_2_ is a conserved positive regulator of most TRP channels^20^. The TRPV5-bound PI(4,5)P_2_ structure revealed directly for the first time how a TRP channel is opened by this important endogenous lipid at the molecular level. Similar binding sites for PI(4,5)P_2_ have been theoretically predicted for TRPV1^26^ and TRPV6^14^ channels using computational modeling and experimental approaches, suggesting that the S4-S5 linker plays an essential role in this interaction. Our cryo-EM structure directly demonstrates an interaction between PI(4,5)P_2_ and TRPV5 through the N-linker, the S4-S5 linker and the S6 helix of each TRPV monomer (Fig. 2). Comparison between the PI(4,5)P_2_ -bound and lipid-bound TRPV5 structures revealed that binding of PI(4,5)P_2_ near to the lower gate of the TRPV5 channel induces conformational changes in the pore, which allows for the flow of Ca^2+^ ions (Fig. 3, Fig. 7).

**Figure 7.**
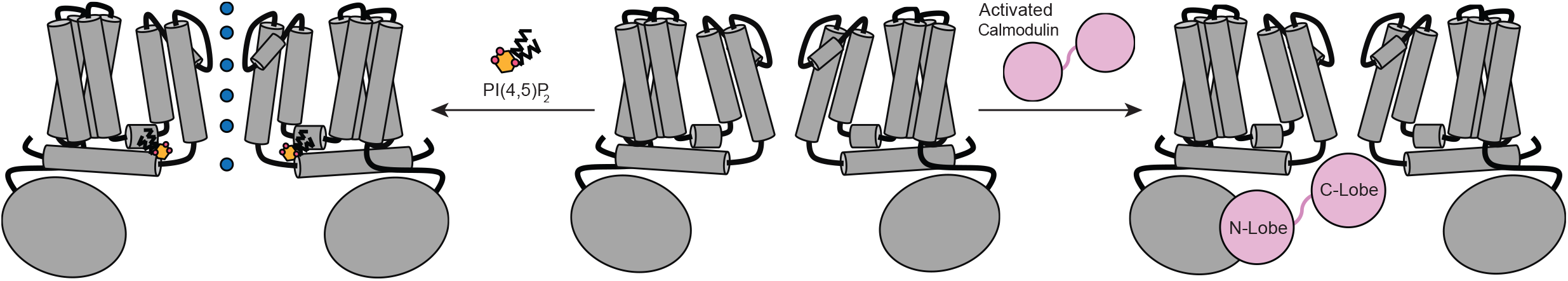
TRPV5 activation by PI(4,5)P2 and inactivation by CaM. A schematic representation of the proposed binding of both PI(4,5)P2 and CaM to TRPV5, inducing either channel activation or inactivation. A dimer of TRPV5 is shown as a grey diagram, PI(4,5)P2 is shown in orange, CaM is shown in pink and the blue circles indicate the flow of calcium ions.

The CaM-bound structure has provided the first structural insight into how TRPV5 ion flow is inhibited by CaM. It is clear that the binding of a single CaM molecule completely obstructs the intracellular side of the pore, effectively blocking ion permeation (Fig. 7). This study also showed that while CaM binds to the predicted areas in the C-terminus of TRPV5, it also directly interacts with the C-terminus at a novel binding site (Fig. 6). Taken together, the lipid- and CaM-bound TRPV5 structures provide additional evidence for the static nature of some TRPV subfamily channels^15,25^. The minimal movement (RMSD ~0.6Å) between the two structures implies that large conformational changes in the upper and lower gates of TRPV5 may not be necessary for effective endogenous inhibition.

It has previously been shown that the N- and C-lobes of CaM have different affinities for calcium ions, with the C-lobe having a 6-fold higher affinity for calcium ions than the N-lobe^27^. Other groups have proposed that this allows the CaM C-lobe to be constantly bound to the distal C-terminus of TRPV5 at normal cellular calcium levels, keeping a constant supply of CaM near the pore^23,28^. In this model, the N-lobe only binds to the C-terminus of TRPV5 when calcium is high and the pore needs to be closed, and the second CaM binding event may stabilize the position of CaM at the base of the pore and lead to the occlusion of the channel^23,28^. The new CaM N-lobe binding site at TRPV5 residues Asn640-Lys652 could compliment the previous model, by potentially “reeling in” CaM when calcium levels are high and bringing the CaM C-lobe into close proximity with the base of the pore.

Overall, our structural studies have provided novel molecular insights into the endogenous modulation of TRPV5 that could guide novel therapeutics design for a variety of calcium dependent kidney diseases.

## METHODS

### Expression and purification of TRPV5

TRPV5 was expressed as reported previously^4^. In short, full-length rabbit TRPV5 was expressed with a 1D4 affinity tag in *S. cerevisiae*^29,30^. The membranes were lysed and harvested using a M-110Y microfluidizer and ultracentrifugation. TRPV5 was purified using previously published methods^4^. Briefly, membranes containing TRPV5 were solubilized in Buffer A (150mM NaCl, 2mM TCEP, 1mM PMSF, 20 mM HEPES pH8, 10% glycerol and 0.87 mM LMNG). Non-soluble material was removed via ultracentrifugation. Detergent solubilized TRPV5 was then purified using CnBr-activated Sepharose 4B beads conjugated to 1D4 specific antibodies. The beads were washed with Buffer B (150mM NaCl, 2mM TCEP, 20 mM HEPES pH8, and 0.064 mM DMNG) and TRPV5 was eluted using Buffer B with the addition of 3 mg/mL 1D4 peptide.

The sample involved in the reconstruction of lipid-bound TRPV5 in detergent was then subjected to size exclusion chromatography (Superose 6, GE Healthcare) in Buffer B. This sample was concentrated to ~2.5mg/mL and incubated with soluble diC_8_ PI(4,5)P_2_ (dioctanoyl Phosphatidylinositol 4,5-bisphosphate) at a final concentration of 200μM for 30 minutes prior to vitrification.

For the sample that was reconstituted into nanodiscs, after elution from the Sepharose beads TRPV5 was incubated with MSP_2_N2 and soy polar lipid extract (Avanti Polar Lipids, Inc.) in a molar ratio of 1:1:200 (TRPV5:MSP_2_N2:Soy Polar Lipids) for 30 min. MSP_2_N2 (Addgene) was expressed and purified as previously published^31^, except the protein was induced at 28°C instead of 37°C. Soy polar lipids were dried under nitrogen flow for 3 hrs prior to reconstitution and dried lipids were resuspended with a 1:200 molar ratio of lipids to DMNG before being added to the protein mixture. Bio-Beads (Bio-Rad, Bio-Beads SM-2 Absorbent Media) were added to the TRPV5, MSP, lipid mixture for 1 hr. Fresh Bio-Beads were then added and allowed to incubate overnight. The reconstituted nanodiscs were further purified using size exclusion chromatography (Superose 6, GE Healthcare) in Buffer C (150mM NaCl, 2mM TCEP and 20 mM HEPES pH 8). This sample was concentrated to ~2.8mg/mL and incubated with soluble diC_8_ PI(4,5)P_2_ at a final concentration of 400μM for 30 minutes prior to vitrification.

The TRPV5 sample used for the reconstruction of CaM-bound TRPV5 was incubated for 1 hr with 10mM CaCl_2_ and purified rat CaM (a gift from the Dr. Zhang lab at Thomas Jefferson University, purified as previously published^32^) at a molar ratio of 1:20 (TRPV5 tetramer:CaM) after elution from the Sepharose beads. The TRPV5-CaM mixture was then further purified using size exclusion chromatography (Superose 6, GE Healthcare) in Buffer B. No chelating agents were added to Buffer B; thus, trace amounts of calcium are present in the final sample. The peak containing CaM-bound TRPV5 was concentrated to ~3mg/mL.

### Cryo-EM data acquisition

For all samples, fluorinated Fos-Choline-8 was added to the concentrated protein to a final concentration of 3mM immediately prior to vitrification. Samples were double blotted on 200 mesh Quantifoil 1.2/1.3 grids (Quantifoil Micro Tools) with 3.5μL per blot and plunge frozen in liquid ethane using a Vitrobot (Thermo Fisher Scientific). Grids containing TRPV5 and diC_8_ PI(4,5)P_2_ in detergent as well as grids with CaM-bound TRPV5 were imaged with a 300kV Titan Krios microscope equipped with a Gatan K2 Summit direct detector camera. Super resolution movies (50 frames) were captured for 10 sec each with one frame collected every 0.2 sec. The resultant pixel size and dose rate were 0.55 Å/pix and ~8 electrons/pix/sec, respectively. Images were collected in a defocus range between 1.0-2.5μm under focus in an automated fashion using Leginon software^33^.

Grids containing TRPV5 and diC_8_ PI(4,5)P_2_ in nanodiscs were also imaged with a 300kV Titan Krios microscope equipped with a Gatan K2 Summit direct detector camera. Super resolution movies (40 frames) were captured with an 8 sec exposure time in super resolution mode resulting in a pixel size and dose rate of 0.532 Å/pix and ~6 electrons/pix/sec, respectively. Images were collected in a defocus range between 1.25-2.5μm under focus.

### Image processing

#### Lipid-bound TRPV5 in detergent

MotionCor2^34^ was used to correct for beam induced motion and to bin the images to a final pixel size of 1.1Å, producing both summed and dose weighted micrographs. CTF estimation of the summed micrographs was preformed using Gctf^35^. All other image processing was performed using RELION on the dose weighted micrographs unless otherwise mentioned^18,19^. Approximately 2,500 particles were picked manually from 3,889 micrographs and sorted into 2D classes. The best classes were used as templates for autopicking. After autopicking, 2D classification was used to remove false positives and suboptimal particles. The remaining ~241,000 particles were reconstructed into a single electron density map with C4 symmetry using the 3D autorefine option in RELION followed by 3D classification into 8 classes without assigning angles to the particles. The initial model used for 3D refinement and classification was produced by applying a low-pass filter of 60Å to the previously published TRPV5 cryo-EM structure^4^. The best class by manual inspection underwent multiple rounds of 3D refinement followed by 3D classification until the best ~26,000 particles were able to be reconstructed into 4.4Å map. The mask used for this reconstruction was created from the original 3D autorefinement of ~241,000 particles adjusted to a threshold of 0.005, lowpass filtered to 15Å and a soft edge of 5 pixels was applied. A separate dataset collected under the same conditions on grids prepared at the same time as the sample above produced 3,062 additional micrographs. The same methods as above were implemented and resulted in a map containing ~19,000 particles that was reconstructed to 4.2Å. These particles were then combined using the JoinStar command in Relion to produce a particle set of ~45,000 particles that were refined to 4.1Å in RELION (Sup. Fig. 2). A b-factor of −174 was then applied to the unsharpened map using Bfactor software and the final resolution of 3.9Å was determined using rmeasure software^36,37^. Local resolutions were estimated using the RESMAP software^38^.

#### PI(4,5)P_2_ bound TRPV5 in nanodiscs

MotionCor2^34^ was used to correct for beam induced motion and to bin the images to a final pixel size of 1.1Å, producing both summed and dose weighted micrographs. CTF estimation of the summed micrographs was preformed using Gctf^35^. All other image processing was performed using RELION on the dose weighted micrographs unless otherwise mentioned^18,19^. To create templates for autopicking, 2,112 particles were manually picked from 1,575 micrographs and sorted into 21 classes. The best classes were used to autopick 493,047 particles which were then subjected to 2D classification into 200 classes to remove false positives and suboptimal particles. The best classes contained 121,980 particles, which were refined to 4.4Å using 3D AutoRefine with C4 symmetry in the absence of a mask using the same initial model as in the lipid-bound structure. A mask was created of this structure by adjusting the threshold to 0.005, lowpass filtering to 15Å and applying a soft edge of 5 pixels. The 4.4Å structure was postprocessed using the above mask to produce the Initial 3D Reconstruction at 4.3Å (Sup. Fig. 4). A cylinder with a radius of 40Å and height of 15Å was created in Chimera, centered on the pore at the height of the TRP helices. This cylinder was used to make a mask at a threshold of 0.5, an extended edge of 2 pixels, and a soft edge of 5 pixels. This mask was then used for particle subtraction, where everything outside of this mask was subtracted, as previously described^39^. The subtracted particles were then sorted by 3D classification into 5 classes without assigning angles, yielding three high resolution classes. Two of these classes had density for PI(4,5)P_2_ (total of ~72,000 particles) and one had no PI(4,5)P_2_ density (~30,000 particles). We swapped both sets of subtracted particles back to the unsubtracted particles and refined both sets using the mask previously used to refine the ~121,000 particle map. The ~30,000 particles for the class without PI(4,5)P_2_ were refined to 4.5Å in RELION. Postprocessing was run on this data in RELION using the same mask as previously with an applied b factor of -200, leading to a final resolution of 4.4Å. The ~72,000 particles were then sorted again by 3D classification into 3 classes without assigning angles. The best of these classes consisted of ~25,500 particles and was refined to 4.1Å in RELION. Postprocessing was run on this data in RELION using the same mask as previously with an applied b factor of -150, leading to a final resolution of 4.0Å (Sup. Fig. 4). Local resolutions were estimated using RESMAP^38^.

#### CaM-bound TRPV5

For the CaM-bound dataset, MotionCorr2^34^ was used without gain correction and without binning to produce both summed and dose weighted micrographs with a final pixel size of 1.1Å. CTF estimation of the summed micrographs was preformed using Gctf^35^. All other image processing was performed using RELION on the dose weighted micrographs unless otherwise mentioned^18,19^. To create templates for autopicking, 3,125 particles were manually picked from 2,118 micrographs and sorted into 30 classes. The best classes were used to autopick 391,648 particles which were then subjected to 2D classification into 100 classes to remove false positives and suboptimal particles. The best classes contained 153,776 particles, which were refined to 5.4Å using 3D AutoRefine with C1 symmetry in the absence of a mask using the same initial model as in the lipid-bound structure. At this point, there was clearly density present in the region between the ankyrin repeat domains of the TRPV5 monomers, but it appeared to be asymmetrical. To make sure that information for all the asymmetrical features was aligned in the same orientation, we decided to use symmetry expansion. The ~154,000 particles from the initial C1 reconstruction were refined in C4 to generate symmetry operators. Relion symmetry expansion in C4 was used to expand the particle stack and ensure that all the information for CaM was present in all four possible orientations. A sphere of radius 25Å was created in Chimera and placed at the base of the pore so that the entire region with density for CaM was in the sphere. A mask was made of this sphere in RELION, at a threshold of 0.1, extended by 5 pixels, and with a soft edge of 5 pixels. This spherical mask was used for particle subtraction, as described above, and then the subtracted particles were sorted into 5 classes with 3D classification. The resulting classes captured CaM in the 4 possible orientations, with approximately equal numbers of particles, as well as a class without CaM bound. The class with the best resolution was chosen, consisting of ~129,000 particles, and we reverted to the unsubtracted particles and refined these particles. There was still some heterogeneity in the CaM, so we further classified the particles into 3 classes, resulting in a class with ~47,500 particles that refined to 4.7Å in RELION. After postprocessing in RELION with an applied b factor of -200, the, this data reached a resolution of 4.4Å (Sup. Fig. 11). Local resolutions were estimated using RESMAP^38^.

### Model Building

The previously published model of TRPV5 bound to econazole (PDB 6B5V)^4^ was used as an initial model and docked to the cryo-EM map of lipid-bound TRPV5. The model was manually adjusted in Coot^40^ and refined with imposed 4 fold NCS using phenix.real_space_refinement^41^. The lipid-bound TRPV5 model was docked into the PI(4,5)P_2_-bound TRPV5 map and manually adjusted and refined as before. The lipid-bound TRPV5 model was docked into the CaM-bound TRPV5 map. To model CaM into the extra density, the N-lobe (residues 6-77) and C-lobe (residues 83-147) of a calcium-bound CaM structure (PDB 1CLL) were separated and independently fit in Chimera to the two lobes of density visible at the base of the pore. A portion of the N-lobe was visible only as broken density, and so was removed from the model. The TRPV5 C-terminal peptide from state 01 of PDB 5OEO was fit into the remaining density near the C-lobe in Chimera. An extension of the C-terminus of TRPV5 chain A was modeled into helical density in the peptide binding pocket of the CaM N-Lobe. This assembly was manually adjusted in Coot and refined without NCS using phenix.real_space_refinement.

The final models were randomized in PHENIX^41^ by 0.5Å and refined against one of the half maps. EMAN2 was used to generate FSC curves between this refined test model and each half map as well as between the final model and the summed map for each model. The pore radii were generated using HOLE^42^. Figures were made in Pymol^43^ and Chimera^44^.

### Xenopus oocyte electrophysiology

Oocyte sacs were extracted from female *Xenopus laevis* frogs (*Xenopus* Express) and digested with 0.2 mg/ml collagenase (Sigma) in OR2 solution (82.5 mM NaCl, 2 mM KCl, 1 mM MgCl_2_, and 5 mM HEPES, pH 7.4) overnight at 18 °C. Oocytes were maintained in OR2 solution plus 1.8 mM CaCl_2_ and 1% penicillin/streptomycin (Mediatech) at 18 °C. cRNA was generated using the mMessage mMachine kit (Thermo Fisher); point mutations were introduced using the Quikchange mutagenesis kit (Agilent Genomics). cRNA (20 ng) was injected using a nanoliter injector system (World Precision Instruments). The W583L mutant of TRPV5 was kept in OR2 with 0.1 mM CaCl_2_, to avoid Ca^2+^ overload and cell damage. The experiments were performed 48-72 h after injection. Purified calmodulin from bovine testes was purchased from Sigma.

Two-electrode voltage clamp (TEVC) measurements were performed as described earlier^14^; briefly oocytes were initially placed in a solution containing 97 mM NaCl, 2 mM KCl, 1 mM MgCl_2_, and 5 mM HEPES, pH 7.4, where TRPV5 or TRPV6 currents are largely blocked by Mg^2+^ and trace amounts of Ca^2+^ in the medium. Monovalent currents were initiated with changing the solution containing 96 mM LiCl, 1 mM EGTA, and 5 mM HEPES, pH 7.4. Currents were recorded with thin-wall inner filament-containing glass pipettes (World Precision Instruments) filled with 3 M KCl in 1% agarose. Currents were recorded with a GeneClamp 500B amplifier (Molecular Devices using a ramp protocol from – 100 to + 100 mV (0.25 mV/ms), immediately preceded by a 100-ms step to – 100 mV, applied once every second; holding potential was 0 mV.

Excised inside-out patch clamp experiments were performed as described earlier^13^ using borosilicate glass pipettes (World Precision Instruments) of 0.8–1.7 megaohm resistance. The electrode pipette solution contained 96 mM LiCl, 1 mM EGTA, and 5 mM HEPES, pH 7.4. After establishing giagohm resistance seals on devitellinized *Xenopus* oocytes, the currents were measured using an Axopath 200B amplifier (Molecular Devices). We used a ramp protocol from -100 mV to 100 mV, performed once a second, immediately preceded by a 100-ms step to -100 mV. The perfusion solution contained 93 mM potassium gluconate, 5 mM HEDTA, 5 mM HEPES, with the pH adjusted to 7.4. To obtain 3 μM free Ca^2+^, 2.5 mM Ca^2+^ gluconate was added. The bath was connected with the ground electrode through an agar bridge.

### TRPV6 structural modeling, docking simulations and molecular dynamics simulations

We generated an almost full-length atomistic model of TRPV6 by performing comparative homology modeling using the structure of the closed state of TRPV1 as a template (PDB:3J5P). We extracted the pairwise sequence alignment between TRPV1 and TRPV6 from the multiple sequence alignment containing about 3,000 sequences^45^. We then generated 288 models using ROSETTA^46^ and discarded the 50 ones with the lowest score. We performed clustering on the remaining 248 structures and found two major clusters differing by the conformation of the S2-S3 segment. Due to the absence of significant structural constraints from the template, we modeled this region of TRPV6 ab initio.

To investigate the binding mode of PI(4,5)P_2_ we considered a model ligand and performed a series of stochastic optimizations of its positions with respect to the TRPV6 structure using the program AUTODOCK^47^. The model ligand is a simplified version PI(4,5)P_2_ that contains the entire head group, the glycerol moiety and part of the acyl chains (up to the third carbon atom). This choice allowed us to explore thoroughly the conformational space by ignoring the large number of degrees of freedom describing the configuration of the flexible acyl chain. Since not all the binding poses of the model ligand correspond to a geometrically viable conformation of PI(4,5)P_2_, we post-processed the output of docking and selected only those binding poses in which the acyl chain is correctly positioned with respect to the lipid bilayer. The conformational search was restricted to a specific region of the channel encompassing the S4-S5 linker, the N-term sections of S1, S3 and S5, the C-term section and S2 and S4, the S2-S3 loop, the N-term section of the TRP box and the adjacent part of the cytoplasmic domain. The region was large enough to contain at least two adjacent subunits to allow for the identification of potential interactions at their interface. For each structure of TRPV6 (20 in total) 1,000 docking experiments were performed using the Lamarckian genetic algorithm allowing for a maximum of 250,000 energy evaluations.

We performed molecular dynamics simulations of the PI(4,5)P_2_-TRPV6 and PI(4,5)P_2_-TRPV1 complexes and of the TRPV6 apo channel (with no bound PI(4,5)P_2_). As initial configurations, we considered the structural models with the highest scores. The channels were embedded into a fully hydrated 1-palmitoyl-2-oleoyl-sn-glycero-3-phosphocholine membrane surrounded by a 150 mM KCl solution. Overall, the system contains more than 300,000 atoms. During the initial equilibration protocol, we gradually released each component of the (water and ions, lipid tails, lipid head groups, protein side chains and protein backbone) system from positional restraints. During the production run, trajectories were collected for 0.7 μs. Simulations were performed at constant temperature and pressure (1 atm, 300K) using the Langevin piston approach using NAMD 2.10^48^. The CHARMM36 force field^49^ was used to describe the protein, the ions and the lipid molecules. For the vdW interactions, we used a cutoff of 11 Å with a switching function between 8 and 11 Å. The long-range component of electrostatic interactions was calculated using the Particle Mesh Ewald approach^50^ using a cutoff for the short-range component of 11 Å. The equations of motion were integrated using the RESPA multiple time-step algorithm^51^, with a time step of 2 fs and long-range interactions calculated every other step.

## DATA AVAILABILITY

The cryo-EM density maps and atomic coordinates of all structures presented in the text are deposited into the Electron Microscopy Data Bank and Protein Data Bank under the access codes shown in Supplementary Table 1. All data is available from the corresponding author upon reasonable request.

## ACKNOWLEDGEMENTS

We thank Dr. Ji-fang Zhang and Dr. AjaySingh Tanwar at Thomas Jefferson University for the gift of purified rat calmodulin. We thank Denice Major for assistance with hybridoma and cell culture at the Department of Ophthalmology and Visual Sciences (supported by the National Institutes of Health Core Grant P30EY11373). We thank Yvonne Gicheru and Dr. Sudha Chakrapani at Case Western Reserve University for training and assistance with MSP_2_N2 expression and purification. We acknowledge the use of instruments at the Electron Imaging Center for NanoMachines supported by NIH (1S10RR23057 and 1S10OD018111), NSF (DBI-1338135) and CNSI at UCLA. We also acknowledge microscopist Carol Bator and the use of instrument at the Penn State Cryo Electron Microscopy Facility (University Park, PA). This research was, in part, supported by the National Cancer Institute’s National Cryo-EM Facility at the Frederick National Laboratory for Cancer Research. We thank Dr. David Lodowski at Case Western Reserve University for help in the early stage of the project. This work was supported by grants from the National Institute of Health (R01GM103899 to V.Y.M.-B., R01GM093290 to T.R. and V.C., U24 GM116792 to Z.H.Z and V.Y.M.-B).

## AUTHORS CONTRIBUTIONS

T.E.T.H. conducted protein purification, cryo-EM sample preparation and cryo-EM data collection; T.E.T.H. and R.A.P. performed all cryo-EM data analysis and interpretation; R.A.P. built and refined all atomic models; A.T.Y. performed all electrophysiology experiments and analyzed the resultant data; M.A.K. performed all TRPV6 modeling, docking and molecular dynamic simulations; E.C.F. assisted T.E.T.H. and R.A.P. in data analysis; K.W.H. assisted T.E.T.H. in cryo-EM data collection; A.S. and S.M. trained and assisted T.E.T.H. in cryo-EM sample preparation and screening; Z.H.Z. supervised cryo-EM data collection; V.C. supervised TRPV6 simulations and the subsequent data interpretation; T.R. supervised the electrophysiology experiments and assisted with data analysis; V.Y.M-B. designed and supervised the execution of all experiments in this manuscript; T.E.T.H., R.A.P., T.R. and V.Y.M.-B. wrote the final version of the manuscript; All authors contributed to and reviewed the final manuscript.

---

**Supplementary Table 1.**
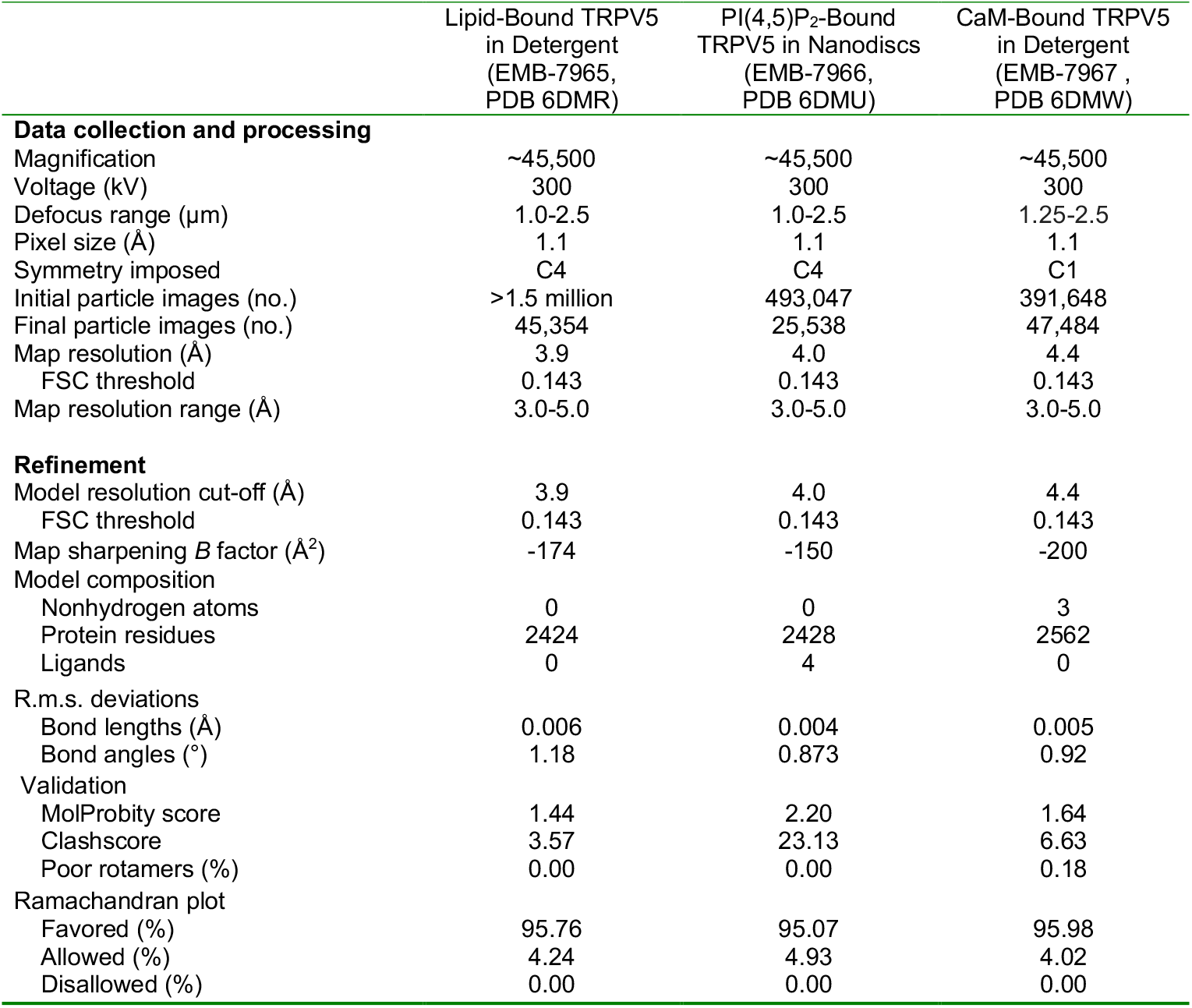
Cryo-EM map details and model statistics.

